# Defining overlooked structures reveals new associations between cortex and cognition in aging and Alzheimer’s disease

**DOI:** 10.1101/2023.06.29.546558

**Authors:** Samira A. Maboudian, Ethan H. Willbrand, William J. Jagust, Kevin S. Weiner, Alzheimer’s Disease Neuroimaging Initiative

**Author notes:** **Corresponding author:** Samira A. Maboudian. Data used in this article were obtained from the Alzheimer’s Disease Neuroimaging Initiative (ADNI) database (adni.loni.usc.edu). ADNI investigators thus contributed to the design and implementation of ADNI and/or provided data but did not participate in analysis or writing of this report. A complete list of ADNI investigators can be found at: http://adni.loni.usc.edu/wp-content/uploads/how_to_apply/ADNI_Acknowledgement_List.pdf.

## Abstract

Recent work suggests that indentations of the cerebral cortex, or sulci, may be uniquely vulnerable to atrophy in aging and Alzheimer’s disease (AD) and that posteromedial cortex (PMC) is particularly vulnerable to atrophy and pathology accumulation. However, these studies did not consider small, shallow, and variable tertiary sulci that are located in association cortices and are often associated with human-specific aspects of cognition. Here, we first manually defined 4,362 PMC sulci in 432 hemispheres in 216 participants. Tertiary sulci showed more age- and AD-related thinning than non-tertiary sulci, with the strongest effects for two newly uncovered tertiary sulci. A model-based approach relating sulcal morphology to cognition identified that a subset of these sulci were most associated with memory and executive function scores in older adults. These findings support the retrogenesis hypothesis linking brain development and aging, and provide new neuroanatomical targets for future studies of aging and AD.

## Introduction

Understanding the relationship between structural brain changes and cognitive decline in aging and neurodegeneration is a fundamental objective of aging research. Empirical findings from large-scale, group-level atrophy studies with hundreds of participants converge on broad, consistent patterns across studies: frontal regions are most affected by atrophy in aging, while temporal regions are more affected in Alzheimer’s disease (AD)^1–5^. However, more specific investigations of individual differences in atrophy patterns are less clear. Measuring changes in sulcal morphology in aging has been proposed as one approach to explore individual differences in cortical structure without the biases introduced from relying on a common stereotaxic space^6, 7^, as well as to provide additional insights about structural change beyond global or regional volume and thickness analyses, since sulcal morphology is thought to reflect underlying white matter architecture and connectivity^8–11^. Such studies in aging have found that sulci are more vulnerable to thinning than gyri^12^ and that sulcal morphology is related to age-related cognitive decline^11, 13^. In AD, sulcal morphology has been proposed as an imaging marker of early-onset AD^14^, has been found to be more diagnostically accurate than traditional measures such as hippocampal volume^6^, and is related to cognitive impairment^6^.

These studies have focused on the largest, deepest sulci across the cortex. However, an emerging area of investigation that has been largely overlooked is the role of tertiary sulci: the smallest, shallowest cortical indentations that emerge latest in development and most recently in primate evolution^9, 15, 16^. Recent research shows that the morphology of tertiary sulci is associated with individual differences in cognitive development^9, 17–19^, as well as with symptoms and onset of disorders including schizophrenia^10^ and frontotemporal dementia^20^. To our knowledge, no prior research has investigated sulcal morphology including tertiary sulci in normal aging or AD.

Here, we build on these findings to investigate tertiary sulci in the aging brain for the first time. One reason tertiary sulci have been overlooked in prior research is that, because of their small size and variability across individuals, they often do not reliably appear in analyses averaged across individuals and are not automatically identified by commonly-used cortical parcellations^9, 15, 21^. Therefore, tertiary sulci must be manually identified by trained raters on each hemisphere, as in previous work^9, 10, 17, 18, 20–27^. Because this process is laborious, these studies of tertiary sulci commonly restrict analyses to a single sulcus or region. Therefore, here we focus on analyzing sulcal morphology in posteromedial cortex (PMC). PMC, which includes the precuneus and posterior cingulate cortex^21, 28, 29^, is vulnerable to atrophy in aging^1, 2^, and age-related alterations in PCC functional connectivity are correlated with cognitive impairment ^30–32^. PMC is also particularly vulnerable to atrophy in AD^4, 33^, is one of the earliest sites of pathological amyloid-β (Aβ) accumulation^34, 35^, and of pathological tau spread beyond medial temporal lobe in AD^36, 37^. Finally, recent work has also clarified the organization of PMC sulci in humans and chimpanzees to include tertiary sulci^21, 22, 38^; however, this modern sulcal model of PMC has yet to be comprehensively applied and examined in aging and AD.

The present study examines changes in PMC sulci in aging and AD by comparing their morphology in young adults, healthy older adults, and older adults with AD. This work was guided by the following questions: (1) Are morphological changes uniform in PMC sulci, or are particular sulci or sulcal types uniquely vulnerable? (2) Is there a relationship between PMC sulcal morphology and cognition in older adults, and is this relationship specific to particular sulci? Importantly, answering these questions is theoretically meaningful. Specifically, the retrogenesis model of aging and AD proposes that brain aging and degeneration mirrors development, first affecting evolutionarily newer and later-developing areas and thus, leading to declines in higher-order cognitive processes^39^. Consequently, we hypothesize that evolutionarily-newer and later-developing tertiary sulci may be particularly vulnerable to aging- or AD-related degeneration and subsequent cognitive decline.

## Results

### Classifying PMC sulcal types in young adults using a data-driven approach

We first manually defined 4,362 PMC sulci in 432 hemispheres in 216 adult participants in three groups: 72 young adults (YA) aged 22-36, 72 cognitively normal older adults (OA) aged 65-90, and 72 older adults with AD aged 65-89 (age-matched to OA). Previous work shows that this sample size is sufficient to quantify reproducible and reliable results relating sulcal morphology to cognition in individual participants ^9, 10, 17–19, 26, 40, 41^. PMC sulci were manually defined on the cortical surface reconstruction of each participant’s T1-weighted MRI scan according to the most recent comprehensive sulcal atlas^38^ (Fig. 1) as in our previous studies^21, 22^ (see Methods for specifics).

**Figure 1.**
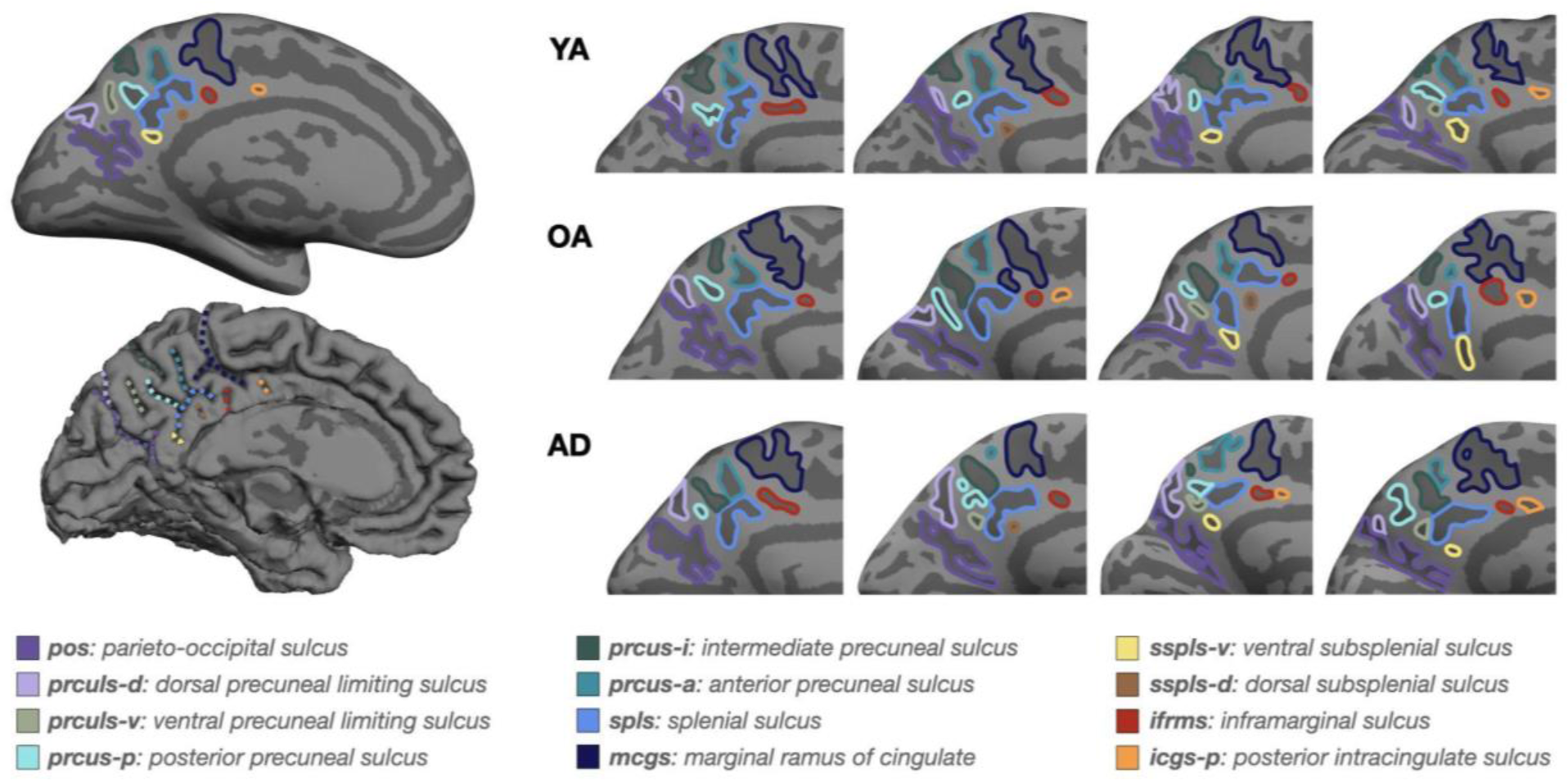
Posteromedial cortex (PMC) sulcal labels and example hemispheres. Left: An inflated (top) and pial (bottom) cortical surface reconstruction of an individual human hemisphere; sulci are dark gray, gyri light gray. Individual PMC sulci are colored according to the legend. Right: 4 example hemispheres within each group, zoomed in to the PMC and depicting variability in sulcal incidence between participants (9-12 sulci per hemisphere: 8 non-tertiary, 1-4 tertiary). Legend: warm colors (right column) are tertiary sulci, cool colors (left) are non-tertiary. YA = Young adult, OA = older adult, AD = older adult with Alzheimer’s.

Sulci are typically categorized as primary, secondary, or tertiary based on their emergence in gestation^42–45^. However, in this study, several sulci included in the modern schematic of PMC sulci^21, 22^ are defined that are not included in this previous work describing gestational sulcation patterns. Therefore, in order to classify sulci independent of these previous classifications, we applied k-means clustering to group sulci based on the primary morphological features considered in this study (sulcal depth and cortical thickness; Fig. 2). In order to limit the effects of age- and AD-related atrophy on sulcal group classification in cases of discrepancies (e.g., a different number of clusters identified across age (OA vs. YA) and hemispheres (OA vs. YA vs. AD)), the YA clustering was used to cluster sulci using this data-driven approach.

**Figure 2.**
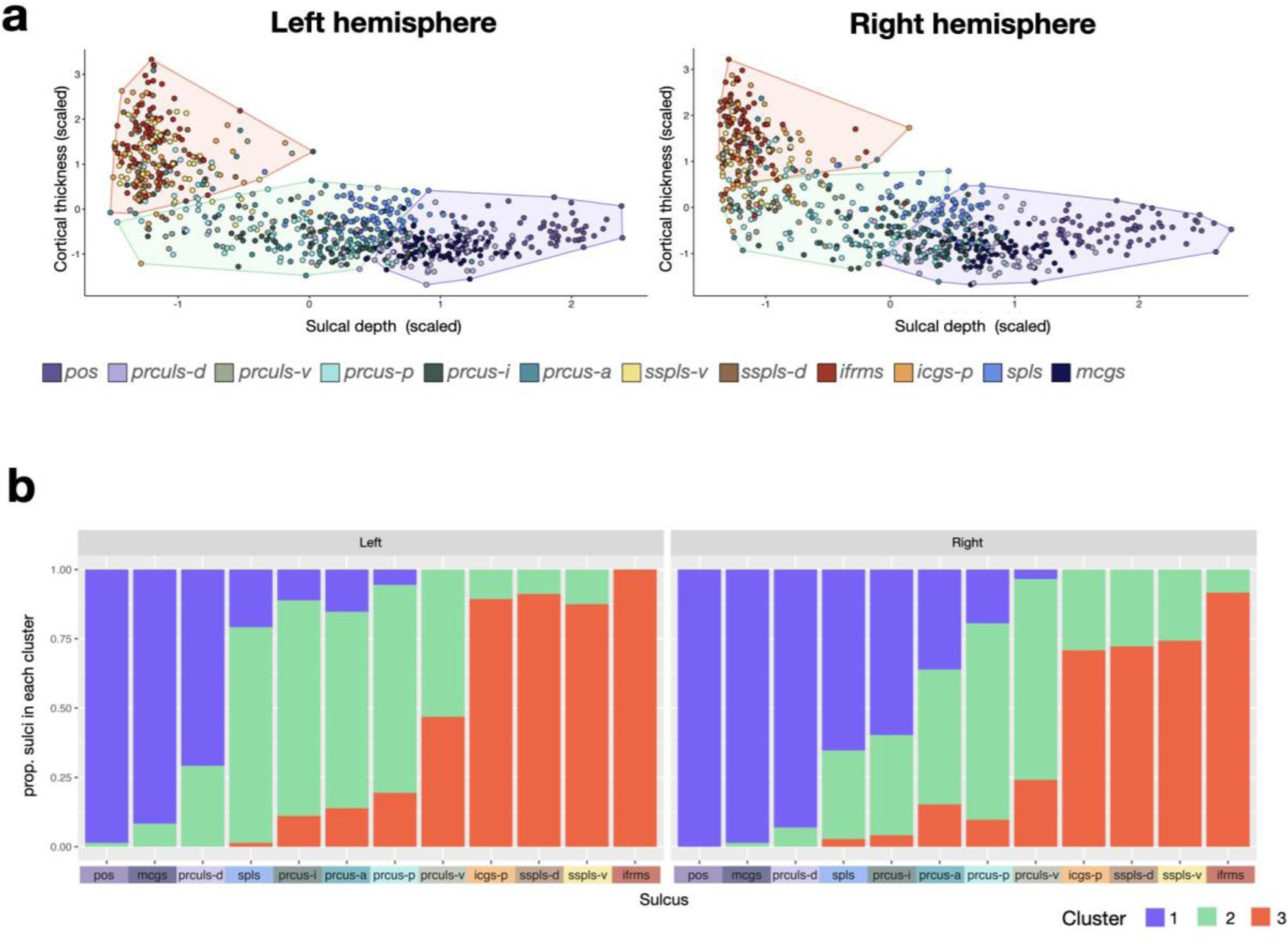
Clustering of sulci into groups by morphological features. **a** Results of k-means clustering in the young adult group, using the *Nbclust* R package to determine the most optimal number of clusters. Each point represents one sulcus (colored by sulcal label), with clustering group overlayed (cluster colors same as in (**b**)). **b** Proportion of observations for each sulcus that occur within each cluster observed in (**a**). The best cluster assignment for each sulcus is indicated by the cluster group that it is most likely to appear in.

The *Nbclust* package in R was used to determine the optimal number of clusters in a data-driven manner blind to sulcal type (Methods). The optimal number of clusters for YA was three. These three clusters corresponded with morphological factors that classically differentiate sulci from one another — such as larger, deeper primary sulci and smaller, shallower tertiary sulci. For example, four small and shallow sulci (ifrms, icgs-p, sspls-d, and sspls-v) are included in one cluster, while the eight other larger and deeper sulci are included in two separate clusters in YA (Fig. 2). Contrary to the stable clustering of the four small and shallow sulci between hemispheres, there was significant variability regarding which large and deep sulci were assigned to cluster 1 or 2 between hemispheres (Fig. 2). As such, when relevant in subsequent analyses, we considered two types of models: a broad model that lumped sulci in two groups (clusters 1 and 2 as “non-tertiary”; cluster 3 as “tertiary”) and a more fine-grained model that considered each sulcus separately.

### PMC tertiary sulci show more age- and AD-related thinning compared to non-tertiary sulci with the strongest effects for newly uncovered sulci (ifrms and sspls-v)

To quantify the effects of sulcal thinning in PMC across groups and hemispheres, we ran a linear mixed-effects model with group (YA, OA, AD), sulcal type (tertiary or non-tertiary), and hemisphere (left or right) as predictors, including all interactions (Fig. 3A). ANOVA F-tests were applied to each model. We observed a main effect of group [F(2,213) = 180.98, *p* < 0.0001, η^2^ = 0.63], sulcal type [F(1,425) = 3491.42, *p* < 0.0001, η^2^_P_ = 0.89, in which tertiary sulci were thicker], and hemisphere [F(1,213) = 4.34, *p* = 0.038, η^2^_P_ = 0.02; in which right hemisphere sulci were thicker], as well as a group x sulcal type interaction [F(2,425) = 37.70, *p* < 0.0001, η^2^ = 0.15]. Post hoc tests revealed that each pairwise group comparison was significant (*p* < 0.0001), with YA having thicker sulci than OA, and OA thicker than AD; each pairwise group comparison was also significant for each sulcal type category separately, but group differences were greater among tertiary sulci (each comparison *p* < 0.0001; Supplementary Table 1).

**Figure 3.**
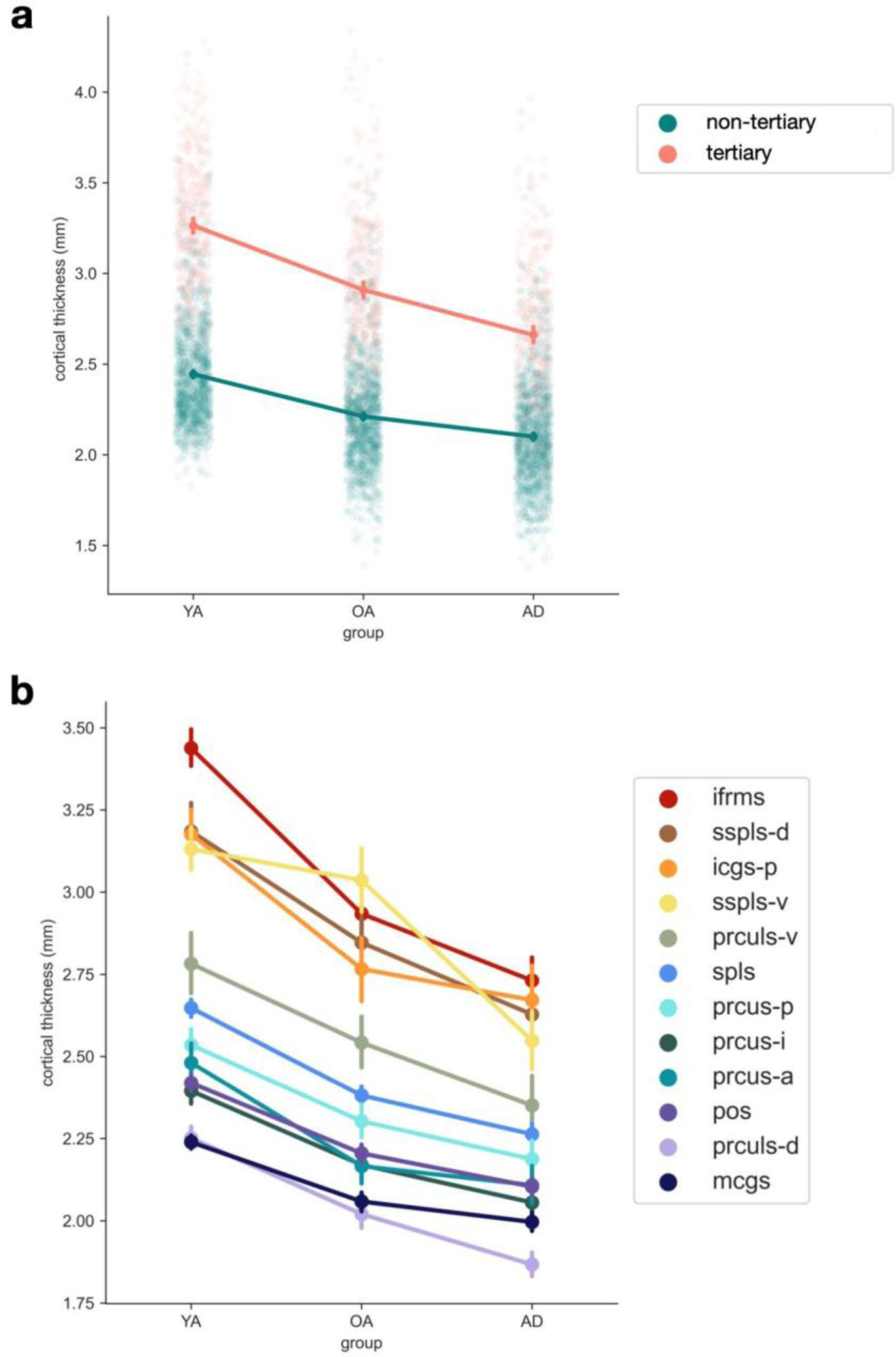
Posteromedial cortex tertiary sulci show more age- and AD-related thinning compared to non-tertiary sulci. **a** Sulcal cortical thickness (in mm) across groups (YA=younger adult, OA=cognitively normal older adult, AD=older adult with Alzheimer’s disease) by sulcal type. Tertiary sulci show greater atrophy across groups than non-tertiary sulci: a linear mixed effects (LME) model with group (YA, OA, AD), sulcal type (tertiary or non-tertiary), hemisphere (left or right) and their interactions as predictors shows a main effect of group (*p<*0.0001), sulcal type (*p<*0.0001), and hemisphere (*p<*0.05), as well as a group x sulcal type interaction (*p<*0.0001). Results of post hoc tests are summarized in Supplementary Table 1A. The points represent the mean thickness, while vertical bars represent the bootstrap 95% CI. **b** Same as (**a**), but showing each sulcus separately, colored according to the legend. An LME model with group (YA, OA, AD), sulcus, hemisphere (left or right) and their interactions as predictors shows a main effect of group (*p<*0.0001), sulcus (*p<*0.0001), and hemisphere (*p<*0.05), as well as a group x sulcus interaction (*p<*0.0001) and a group x hemisphere interaction (*p<*0.05). Results of post hoc tests are summarized in Supplementary Table 1B.

In order to investigate sulcus-specific trends, we also ran the same model but with sulcus as a predictor instead of sulcal type (Fig. 3B). We observed a main effect of group [F(2,213) = 177.31, *p* <0.0001, η^2^ = 0.62], sulcus [F(11,3864) = 560.02, *p* < 0.0001, η^2^ = 0.61], and hemisphere [F(1,213) = 6.60, *p* = 0.011, η^2^ = 0.03; in which right hemisphere sulci were thicker], as well as group x sulcus [F(22,3856) = 10.41, *p* < 0.0001, η^2^ = 0.06] and group x hemisphere interactions [F(2,213) = 3.67, *p* = 0.027, η^2^ = 0.03]. Post hoc tests revealed that each pairwise group comparison was significant (*p* < 0.0001), with YA having thicker sulci than OA, and OA thicker than AD; each pairwise group comparison was also significant for each hemisphere separately (all *p* < 0.0001), but group differences were slightly greater in the right hemisphere for the YA-OA difference and in the left hemisphere for the CN-AD difference. Within each sulcus, pairwise group comparisons were significant (*p* < 0.05), except for icgs-p (*p*=0.1898), mcgs (*p*=0.2441) and prcus-a (*p*=0.3364) for the AD-OA comparison, and sspls-v for the OA-YA comparison (*p*=0.0809) (see Supplementary Table 1 for all *p*-values). The strongest effects occurred in tertiary sulci: the ifrms for the OA-YA comparison and the sspls-v for the AD-OA comparison (Supplementary Table 1). Interestingly, the ifrms was recently shown to be the locus of the thickest portion in PMC^21^, and the sspls-v was just recently uncovered^22^.

As the ifrms is situated anteriorly in PMC, we tested the hypothesis that there is a relationship between the mean anterior coordinate of each sulcus (using the FreeSurfer RAS coordinate system; Methods) and the amount of atrophy in that sulcus across groups (Supplementary Fig. 1). This quantification showed that the amount of sulcal atrophy was correlated with the anterior coordinate when comparing sulcal cortical thickness differences in aging (between YA and OA; r^2^ = 0.43; *p* < 0.03), but not when comparing OA and AD (r^2^ = 0.0; *p* > 0.9). Instead, in AD, there is relatively less atrophy in more anterior sulci and more atrophy in posterior sulci (Supplementary Fig. 1).

As sulcal depth has been reported to decrease in aging, we ran the same linear mixed-effects model described above for sulcal depth instead of cortical thickness (Supplementary Fig. 2) especially because previous results have focused on primary sulci and have shown mixed results that vary by sulcus (with some sulci showing no decrease in depth with age)^46–49^. We observed a significant effect of group [F(2,213) = 3.234, *p* = 0.041, η^2^ = 0.03], sulcal type [F(1,425) = 3242.19, *p* <.0001, η^2^ = 0.88; in which tertiary sulci were shallower], and hemisphere [F(1,213) = 20.443, *p* <.0001, η^2^ = 0.09; in which right hemisphere sulci were shallower], but no significant interactions. While the group effect is driven by a difference between YA relative to OA depth, this effect does not survive multiple comparisons correction (*p* = 0.065); indeed, post hoc pairwise comparisons between all 3 groups showed no significant pairwise differences (all *p* > 0.05).

### Recently identified tertiary sulci are most associated with cognition in older adults

To investigate the relationship between PMC sulcal thinning and cognitive decline, we employed a data-driven pipeline described previously^9, 18, 19^. We first implemented a LASSO regression model predicting ADNI memory composite (ADNI-Mem) scores using the sulcal cortical thickness of the 9 most common PMC sulci, split by hemisphere (Methods). LASSO regression performs feature selection by shrinking model coefficients and removing the lowest from the model, with the sulci that are the strongest predictors of ADNI-Mem scores remaining in the final model. We used leave-one-out cross-validation to determine the value of the LASSO shrinking parameter (α), iteratively fitting models with different α values and choosing the one that minimizes cross-validated mean squared error. This procedure indicates that the thickness of a subset of PMC sulci in each hemisphere is most strongly related to ADNI-Mem scores: pos, prcus-p, spls, and sspls-v in the left hemisphere and prculs-d, prcus-p, ifrms, and sspls-v in the right (Fig. 4A). Intriguingly, this model identifies recently uncovered tertiary sulci (ifrms; sspls-v)^21, 22^ as explaining a significant amount of variance. Because not all models examined are nested, we compared model performance with the Akaike Information Criterion (AIC). Lower AIC is better, and ΔAIC > 2 suggests an interpretable difference between models, while ΔAIC > 10 suggests a substantial difference between models.

**Figure 4.**
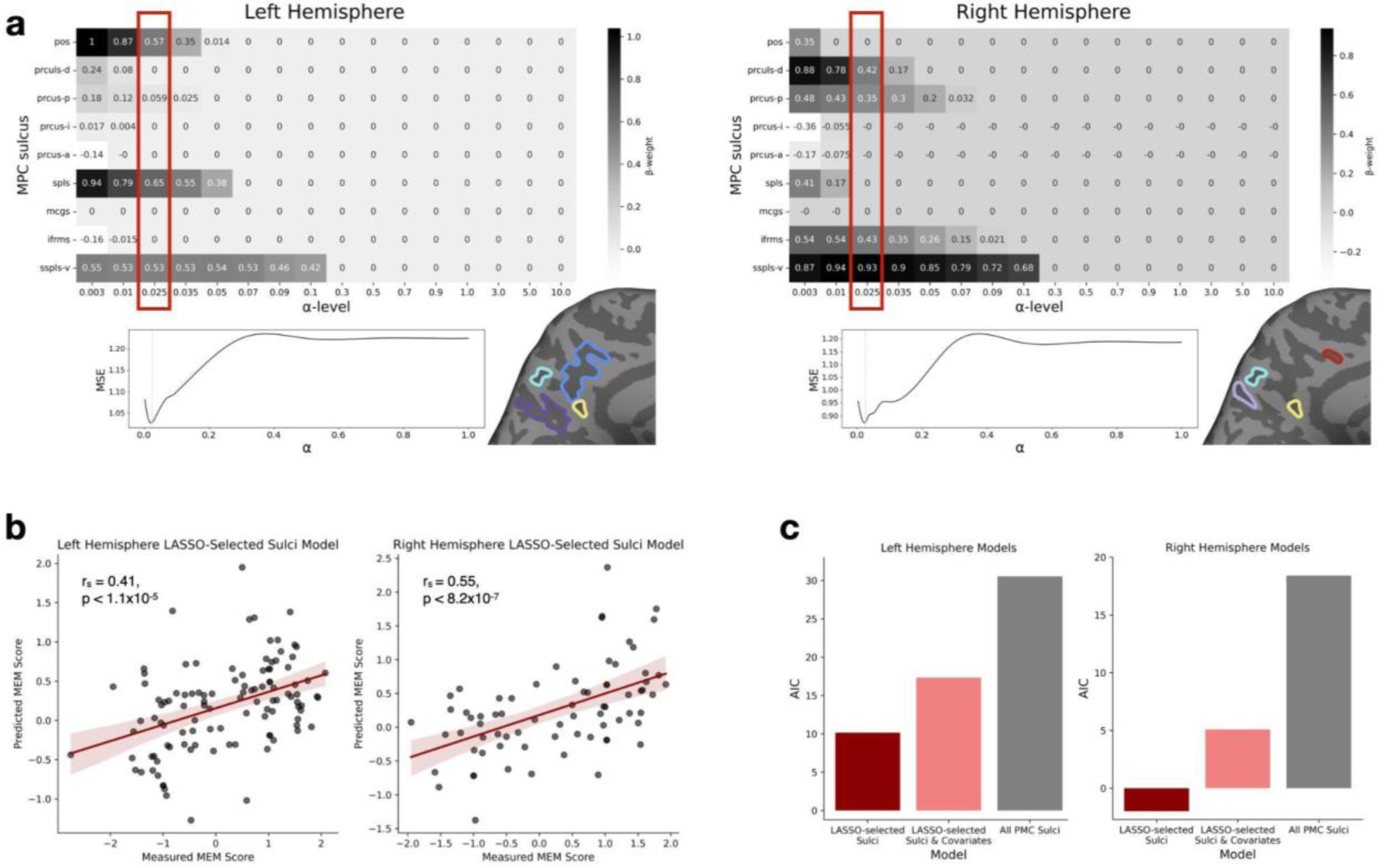
A subset of posteromedial cortex (PMC) sulci are most associated with memory scores. **a** LASSO regression results for models predicting ADNI Memory composite (ADNI-Mem) scores from left hemisphere (left) and right hemisphere (right) PMC sulcal thickness. Top: Beta coefficients for each predictor (sulcal cortical thickness) over a range of shrinking parameter (α) values. Red box depicts values for the chosen model (α value that minimizes MSE_CV_). Bottom left: MSE_CV_ at each α level; α was selected to minimize MSE_CV_ (dotted line). Bottom right: example participant PMC with model-selected sulci highlighted. **b** Spearman’s correlation (r_s_) between actual ADNI-Mem scores and predicted scores from the LOOCV for the best performing LASSO-selected model in the left (left plot) and right (right plot) hemispheres, which had the predictors indicated by the red boxes in (**a**). **c** Model comparison (Akaike information criterion, AIC) of the model with the thickness of sulci selected by the LASSO regression as predictors; a model with the thickness of LASSO-selected sulci and age, sex, and education as predictors; and a model with the thickness of all PMC sulci as predictors for left and right hemispheres. LASSO-selected models have a lower AIC in both hemispheres compared to other models.

We then assessed the relationship between LASSO-selected sulci and ADNI-Mem scores using a linear regression model. For each hemisphere, we compared a model with the thickness of all sulci to a model with the thickness of LASSO-selected sulci (for that hemisphere) only. In the right hemisphere, for the LASSO-selected sulci model, there was a significant association between predicted and actual ADNI-Mem scores (r_s_ = 0.55, *p* < 8.2×10^−7^; Fig. 4B), and the model AIC was −1.99 (Fig. 4C). This model outperformed a model with the thickness of all PMC sulci in the right hemisphere (AIC = 18.40; Fig. 4C). In the left hemisphere, for the LASSO-selected sulci model, there was also a significant relationship between predicted and actual ADNI-Mem scores (r_s_ = 0.41, *p* < 1.1×10^−5^; Fig. 4B), and the model AIC was 10.17 (Fig. 4C). This model outperformed a model with the thickness of all PMC sulci in the left hemisphere (AIC = 30.54; Fig. 4C). ΔAIC between the LASSO-selected right hemisphere sulci thickness model and the LASSO-selected left hemisphere sulci model for ADNI-Mem scores is −12.15, indicating the thickness of right hemisphere LASSO-selected sulci is strongly preferred as a predictor of ADNI-Mem scores.

Available covariates (age, education, and sex) were not associated with ADNI-Mem scores (age r_p_ = −0.033, *p* > 0.6; education r_p_ = 0.12, *p* > 0.14; sex *p* > 0.14); unsurprisingly, including them in the model did not improve model AIC in either hemisphere (Fig. 4C). Further, even though there were no significant depth effects across groups, in order to determine whether these relationships apply to other morphological measures beyond thickness, we also assessed the relationship between the depth of the same LASSO-selected sulci in each hemisphere and ADNI-Mem scores. The LASSO-selected sulcal depth model predictions were not correlated with actual ADNI-Mem scores in either hemisphere, and the thickness model was substantially preferred over the depth model in both hemispheres (Supplementary Fig. 3).

In order to determine whether the morphology of these sulci is related to other cognitive domains beyond memory, we also assessed the relationship between the same LASSO-selected sulci in each hemisphere and ADNI executive function (ADNI-EF) scores. In the right hemisphere, for the LASSO-selected sulci model, there was a significant association between predicted and actual ADNI-EF scores (r_s_ = 0.48, *p* < 2.2×10^−5^; Fig. 5A), though this relationship is weaker than with ADNI-Mem scores, and the model AIC was 22.43 (Fig. 5B). We compared model performance between the LASSO-selected sulcal thickness models for ADNI-Mem and ADNI-EF: ΔAIC = −24.41, indicating that the ADNI-Mem model is substantially preferred (Fig. 5B). In the left hemisphere, for the LASSO-selected sulci thickness model, there was also a significant relationship between predicted and actual ADNI-EF scores (r_s_ = 0.40, *p* < 2.8×10^−5^; Fig. 5A), though again this relationship is weaker than with ADNI-Mem scores, and the AIC was 56.29 (Fig. 5B). The ΔAIC between this model and the left hemisphere ADNI-Mem thickness model (ΔAIC = −46.13) indicates that in the left hemisphere the ADNI-Mem model is again substantially preferred over the ADNI-EF model (Fig. 5B).

**Figure 5.**
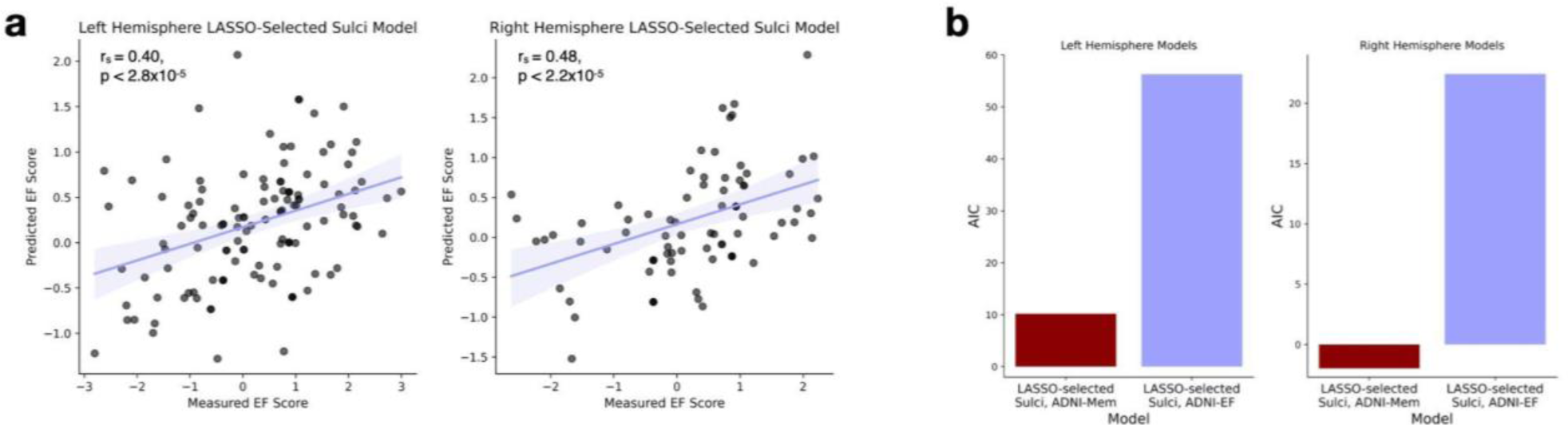
The thickness of LASSO-selected sulci in both hemispheres are associated with ADNI-EF scores. **a** Spearman’s correlation (r_s_) between actual and predicted ADNI Executive Function composite (ADNI-EF) scores from a linear regression model using LASSO-selected PMC sulcal thickness to predict ADNI-EF scores instead of Memory Composite (ADNI-Mem) scores (as in Fig. 4), for both hemispheres. **b** Model comparison (AIC) of LASSO-selected sulci thickness models predicting ADNI-EF scores and ADNI-Mem scores (left: left hemisphere sulcal thickness models, right: right hemisphere models); ADNI-Mem thickness models have a substantially lower AIC in both cases than ADNI-EF models.

## Discussion

In order to investigate and detect potential changes in sulcal morphology in aging and AD at a previously unexplored level of neuroanatomical detail in PMC — including recently uncovered sulci presently excluded from neuroanatomical atlases and neuroimaging software packages — we manually defined 4,362 PMC sulci across 432 hemispheres. Implementing a data-driven approach to classify sulcal types based on multiple morphological features revealed three groupings in young adults, but fewer in older adults. Using these groupings, PMC tertiary sulci (the smallest and shallowest sulci identified by our data-driven approach) show more age- and AD-related thinning compared to non-tertiary sulci (the sulci in the other two groups identified by our data-driven approach), with the strongest effects for two newly uncovered sulci (ifrms and sspls-v). Interestingly, there was an anterior bias for atrophy in aging that is absent in AD. A model-based approach relating sulcal morphology to cognition identified that the thickness of a subset of these PMC sulci (including the two newly identified tertiary sulci) are most associated with memory and executive function scores in older adults, especially in the right hemisphere. Here, we discuss these findings in the context of highlighting (i) the role of tertiary sulci in theories of aging and cognitive development, (ii) the importance of individual-level investigations relating brain structure and cognition, (iii) structural brain changes differentiating aging and AD, and (iv) general limitations and future directions of this study.

The findings of this study relate to classical theories of brain aging and development. The retrogenesis or “last in, first out” model of aging proposes that brain aging mirrors development, with the latest-developing and evolutionarily-newest brain structures (such as prefrontal cortex and association cortices more generally) being the first to degenerate^39^. This theory has also been applied to cognition, with the observation that later-developing cognitive functions (such as executive function) are the first to decline with age. In line with this hypothesis, we find that in PMC, tertiary sulci show the most atrophy in both aging and AD, suggesting that tertiary sulci provide structural evidence in support of this theory. In addition, a classic neuroanatomical hypothesis proposed that morphological changes in tertiary sulci are likely to be associated with the development of higher-order cognitive skills^50^. A growing body of evidence, including the present findings, across age groups, species, and clinical populations supports this theory^9, 10, 17–20, 51^. Combining this theory with the retrogenesis model suggests that these tertiary-prominent morphological changes may also be related to cognitive decline in aging. Our findings support this link, as the tertiary sulci that show the greatest age- and AD-related atrophy are among the sulci with the strongest relationships to cognitive decline in PMC.

Investigating sulcal morphology in aging and AD at the individual level has distinct advantages. For example, applications of the retrogenesis hypothesis to global (e.g., whole brain) cortical atrophy and demyelination patterns have provided mixed results^52, 53^. Our results suggest that investigating brain structure on a finer anatomical scale may provide additional insights not captured by these previous studies. Furthermore, other studies investigating large-scale brain changes in sulci in aging have found that sulci are particularly vulnerable to atrophy in aging compared to gyri^12^ and that sulcal morphology may relate to cognitive decline in aging and AD^6, 11, 13^, but these investigations have all focused on global sulcal morphology or the largest, most prominent sulci across the cortex. Substantial individual variability (including many tertiary sulci) is lost when relying on averaged templates and large group comparisons^6, 9, 15^, suggesting investigations of sulcal morphology at the individual level can provide unprecedented insight into changes in brain structure, given that they appear to show unique atrophy patterns. This is the first study to focus on all sulci—tertiary included—in individual hemispheres across ages and AD in a cortical region, uncovering unique relationships among sulcal types and specific sulci. Further extending these methods to additional areas of the brain in future studies is an intriguing avenue to examine additional individual-level structural changes that may relate to cognitive decline and AD, as well as correlations of sulcal changes in aging and AD between lobes at the level of individually defined sulci.

An additional topic of future study is to further understand the hemispheric differences observed in the present study. For example, the number of observed clusters differed between OA and YA in the right hemisphere only, and right hemisphere sulcal thickness was most associated with cognition (ADNI-Mem and ADNI-EF scores). While global hemispheric differences in atrophy in aging and AD are not established, these findings in PMC are in line with previous work reporting lower regional cortical volume between MCI/AD and controls in right (but not left) precuneus and parietal regions, and lower cortical thickness in right isthmus cingulate in AD compared to MCI^6^. Future work may examine whether these hemispheric differences in sulcal morphology and atrophy are consistent across the cortex, or if they are specific to PMC.

In addition, analyzing sulcal morphology in aging and AD can also provide insight into mechanistic links between structural brain changes and cognitive decline. Previously-reported links between the morphology of tertiary sulci and the development of cognitive skills have been proposed to reflect differences in neural efficiency due to underlying white matter connectivity differences^9, 10^. In aging, sulcal morphology changes (such as sulcal widening and depth) in the most prominent sulci across the cortex have been associated with decreases in white matter volume^54, 55^. These findings have not been extended to include tertiary sulci, which future research could also explore.

Relatedly, improving our understanding of structural changes in the aging brain could aid in differentiating atrophy patterns in healthy aging and AD. Prior work has found that the morphology (including sulcal cortical thickness) of several large, prominent sulci is more accurate to predict AD diagnosis than other more traditional MRI metrics such as hippocampal volume, cortical thickness, and regional volume (of several regions, including precuneus and posterior cingulate gyri)^6^. Including tertiary sulci could further improve the accuracy of such predictive models. Additionally, specific tertiary sulci have been found to have translational applications; for example, the absence of the variable paracingulate sulcus was associated with a significant reduction of age of onset of behavioral variant frontotemporal dementia^20^, and reduced paracingulate sulcal length is associated with increased likelihood of hallucinations in schizophrenia^10^. Investigating the diagnostic potential of these variable tertiary sulci is thus another intriguing avenue for future work, as is investigating relationships between their morphology and pathology. Further, postmortem histological studies suggest that Aβ accumulates in sulci more than gyri^56, 57^, but these findings have not been investigated in tertiary sulci or with in-vivo imaging methods such as Aβ PET imaging, which is another potential avenue of future research.

The main limitation of the present study is the time-consuming process of manually labeling sulci in individual participant hemispheres, which limits the feasible sample size. 4,362 sulci in 432 hemispheres across three groups is a large sample size for an anatomico-cognitive study, but a relatively small sample size compared to studies investigating group-level, cortex-wide structural changes in aging that average across hundreds of participants. Ongoing work is already underway to develop deep learning algorithms to accurately define tertiary sulci automatically in individual participants in lateral prefrontal cortex (LPFC)^58^, posteromedial cortex^21^, and the whole brain^59^. In addition, the age- and AD-related atrophy patterns reported here are cross-sectional. Future studies are underway to extend the cross-sectional findings reported here to longitudinal investigations of sulcal atrophy. Recent results showing a relationship between longitudinal morphological changes in LPFC sulci (including tertiary) and longitudinal changes in cognition in individual participants shows promise for this ongoing and future endeavors^19^.

In summary, we manually identified PMC sulci in individual participant hemispheres and then used a data-driven approach to group sulci by type. Using these groupings, we demonstrated that tertiary sulci in this region atrophy more than non-tertiary sulci in aging and AD, and that a subset of these sulci are also associated with cognitive decline. We connect these findings to theoretical explanations of brain aging and highlight the importance of extending anatomical investigations to include individual-level sulcal analyses. These results bolster mechanistic and theoretical insights regarding structural changes in brain aging, and serve as a foundation for future work further examining unique properties of tertiary sulci in aging and AD.

## Materials and Methods

### Participants

#### Young adults

Data for the young adult cohort analyzed in this study are from the Human Connectome Project Young Adult (HCP-YA) database (https://www.humanconnectome.org/study/hcp-young-adult/overview)^60^. We used data from 72 randomly-selected participants (36 female; age 22-36, µ=29.1 years; Supplementary Table 2). HCP consortium data were previously acquired using protocols approved by the Washington University Institutional Review Board. In this study, we have used the same participants as in our prior work delineating sulcal morphology in human and chimpanzee PMC^22^.

#### Older adults

Data for the older adult cohorts analyzed in this study are from the Alzheimer’s Disease Neuroimaging Initiative (ADNI) (https://adni.loni.usc.edu/). We used MRI scans from 144 older adults: 72 classified as cognitively normal older adults (OA) and 72 with a diagnosis of AD. The 72 OA participants were randomly chosen from the subset of cognitively normal older adults in ADNI (38 female; age 65-90, µ=76.0 years; Supplementary Table 2). The 72 AD participants were chosen from the subset of adults with AD in ADNI who had an MRI and amyloid (Aβ) PET scan within the same year, such that the OA and AD groups were age-matched (35 female; age 65-89, µ=76.2 years; Supplementary Table 2). All AD participants were confirmed to be Aβ positive based on ADNI’s Aβ PET quantification positivity threshold.

### Data Acquisition

#### MRI - Young adults

Anatomical T1-weighted MRI scans (0.8 mm voxel resolution) were obtained in native space from the HCP database, as were outputs from the HCP-modified FreeSurfer pipeline (v5.3.0)^61–64^. Additional details on image acquisition parameters and HCP-specific image processing can be found in Glasser et al.^64^

#### MRI - Older adults

Anatomical T1-weighted MPRAGE anatomical scans were obtained from the ADNI online repository (http://adni.loni.usc.edu). The resolution and exact scanning parameters varied slightly across the sample (see Supplementary Tables 3 and 4 for scanning parameters). Each scan was visually inspected for scanner artifacts, and then cortical surface reconstructions were generated for each participant using the standard FreeSurfer processing pipeline [FreeSurfer (v6.0.0): surfer.nmr.mgh.harvard.edu]^61, 62^.

#### Cognitive data

Cognitive data was obtained from ADNI for both older adult groups. We used the ADNI-Mem memory composite score from the ADNI neuropsychological battery^65^ as a measure of memory performance, and ADNI-EF executive function composite score as a measure of executive function^66^.

### Manual sulcal labeling of PMC sulci

For both cohorts, each cortical reconstruction obtained by the steps above was inspected for segmentation errors (which were manually corrected if necessary). All sulcal labeling and extraction of anatomical metrics were calculated from the FreeSurfer cortical surface reconstructions of individual participants.

As in our prior work^21, 22^, we defined PMC sulci for each participant based on the most recent schematics of sulcal patterning by Petrides^38^ as well as their own pial, inflated, and smoothed white matter (smoothwm) FreeSurfer surfaces. In some cases, the precise start or end point of a sulcus can be difficult to determine on a surface^59^; examining consensus across multiple surfaces allowed us to clearly determine each sulcal boundary in each individual. Sulci were defined in *tksurfer*, where manual lines were drawn on each participant’s FreeSurfer inflated cortical surface to distinguish sulci. For each hemisphere, the location of PMC sulci was identified by trained raters (S.A.M., E.H.W.) and confirmed by a trained neuroanatomist (K.S.W.). See Supplementary Figs. 4 and 5 for all sulcal labels on all older adult hemispheres, and our prior work for all young adult labels^22^.

For this process, we started with the largest and deepest sulci that bound our region of interest (PMC): the parieto-occipital sulcus (pos) posteriorly and the marginal ramus of the cingulate sulcus (mcgs) anteriorly. Next, the splenial sulcus (spls) separates two commonly-delineated subregions of PMC, the precuneus (PrC) superiorly and the posterior cingulate cortex (PCC) inferiorly^21, 22, 67^. PrC contains four consistent sulci: the dorsal precuneal limiting sulcus (prculs-d) and three precuneal sulci (posterior: prcus-p, intermediate: prcus-i, anterior: prcus-a). PCC contains one consistent sulcus: the inframarginal sulcus (ifrms). In addition to these 8 consistent sulci in PMC, there are 4 variably-present sulcal indentations: the ventral precuneal limiting sulcus (prculs-v) in PrC, the posterior intracingulate sulcus (icgs-p) in PCC anterior to the ifrms, and the dorsal and ventral subsplenial sulci (sspls-d and sspls-v, respectively) inferior to the spls. For more detail, see our prior work characterizing PMC sulcal morphology^22^ and Fig. 1 for example participant hemispheres with PMC sulci defined.

### Clustering PMC sulci based on morphology

We used k-means clustering to categorize PMC sulci^19^ as tertiary or non-tertiary based on the morphological metrics considered in this study (sulcal cortical thickness and depth). This analysis was run on scaled morphological data, separately for each group and each hemisphere, and was blind to sulcal type and age. Rather than choosing the number of clusters (k) ourselves, we quantitatively determined the optimal number of clusters in a data-driven manner using the *NbClust* function of the *NbClust* R package — this function uses 30 indices to determine the best number of clusters for the data based on the majority of these indices^68^. K-means clustering was then implemented (with the optimal number of clusters suggested by the *NbClust* results) using the *kmeans* R function.

### Quantification of sulcal morphology

For this study, we extracted and analyzed the cortical thickness and depth of each PMC sulcus, as these are two defining morphological features of cortical sulci that are also of interest in aging^46–49, 69–72^. Though prior work examining sulcal morphology in aging has also studied sulcal width (e.g. ^6, 11, 13, 14, 55^), we found that the current toolbox that extracts this information from individual sulcal labels in FreeSurfer^47^ failed to obtain the width of many smaller sulci (previous studies only examine large sulci). Therefore, we were not able to examine sulcal width in the present study.

#### Cortical thickness

Cortical thickness (in millimeters) was generated for each sulcus using the *mris_anatomical_stats* function in FreeSurfer^61, 62^. Raw (un-normalized) cortical thickness values (in millimeters) are used in the present study as previous studies showed that raw, un-normalized values outperform normalized measures of thickness in MCI and AD^73^.

#### Depth

Sulcal depth was calculated from the native cortical surface reconstruction, measuring from the sulcal fundus to the smoothed outer pial surface. Raw values for sulcal depth (in millimeters) were calculated using a modified version of an algorithm for robust morphological statistics that builds on the FreeSurfer pipeline^47^.

### Morphological comparisons

To compare the cortical thickness of PMC sulci between groups, we ran a linear mixed-effects model (LME) with the following predictors: group (YA, OA, AD), sulcal type (tertiary or non-tertiary), and hemisphere (left or right), and their interactions. Group, hemisphere, and sulcal type were considered fixed effects, and sulcal type was nested within the hemisphere, which was nested within subjects. ANOVA F-tests were applied to each model. We then ran a similar model with the specific sulcal label instead of the sulcal type categories. We also repeated these analyses with sulcal depth instead of cortical thickness. For all post hoc comparisons conducted, *p*-values were corrected with Tukey’s methods, and the Tukey’s-corrected *p* is reported.

### Relating sulcal morphology to cognition

To determine which sulcal measures were most associated with cognition, we employed a least absolute shrinkage and selection operator (LASSO) regression, as described previously^9, 18^. We utilize LASSO regression as a form of feature selection to determine whether the cortical thickness of any PMC sulci in particular predicts memory (ADNI-Mem) scores. We employ this method separately for each hemisphere. LASSO regression requires all individuals to have the same predictors; therefore, in order to include more than one tertiary sulcus while maximizing the available sample size, we included the cortical thickness of all 8 consistent sulci plus the next most consistent PMC sulcus (sspls-v). This resulted in 106 individuals with all 9 of these sulci in the left hemisphere and 70 with all 9 in the right for the LASSO regressions. OA and AD participants were combined to increase the range of scores and sample size available (Supplementary Fig. 6). For the ADNI-EF analyses, one participant with all 9 sulci in the left hemisphere was excluded from analyses because they were missing the ADNI-EF composite score.

This approach allows for data-driven variable selection: LASSO algorithms penalize model complexity by applying a shrinking parameter (α) to the absolute magnitudes of the coefficients, dropping the lowest from the model. LASSO regression increases model generalizability by providing a sparse solution that reduces coefficient values and decreases variance in the model, without increasing bias^74^. To choose the optimal value for α, we use the *GridSearchCV* function of the *Scikit-Learn* Python module to perform an exhaustive search over a range of values, selecting the value that minimizes the cross-validated mean squared error (MSE_CV_) using the following formula:

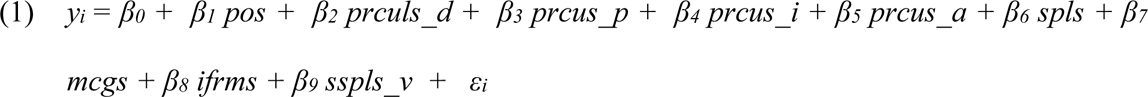

From the LASSO regression with the optimal α value, we identify our model of interest, which best explains memory scores from a subset of the original predictors. One model was identified per hemisphere:

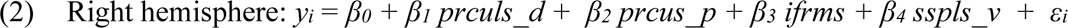

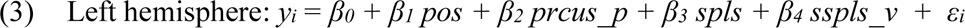

We use linear regression with leave-one-out cross-validation to fit these selected models; all regression models were implemented with the *Scikit-Learn* module. To verify that the results of our feature selection ([2] and [3]) outperform a full model containing all PMC sulci ([1]), we compare the simplified models of interest with the full model, separately for each hemisphere. To determine model specificity, we investigated whether the observed relationship between sulci and task performance generalized to other morphological features or cognitive domains: we compared the original models to models predicting ADNI-EF scores instead of ADNI-Mem scores from sulcal thickness, and to models using the depth of selected sulci as the predictors instead of cortical thickness. We compared the fits of all models of interest using the Akaike Information Criterion (AIC). An AIC>2 suggests an interpretable difference between models, and AIC>10 suggests a strong difference between models (lower AIC suggests better model fit).

### Associations between covariates and cognitive scores

To determine if available covariates (age, education, and sex, summarized in Supplementary Table 2) were associated with ADNI-Mem scores, we calculated Person correlations (r_p_) between age or education and ADNI-Mem scores, and t-tests comparing ADNI-Mem scores between male and female participants.

### Packages used for statistical analyses

All statistical tests were implemented in Python (v3.9.13) and R (v4.0.1). The *Nbclust* package in R was used to determine the most optimal number of sulcal clusters based on morphology, and the R function *kmeans* (from the *stats* R package) was used to perform k-means clustering using this optimal number of clusters. For morphological comparisons, LMEs were implemented with the *lme* function from the *nlme* R package. ANOVA F-tests were implemented with the *anova* function from the built-in *stats* R package, and the effect sizes for ANOVA effects are reported as partial eta-squared (η^2^) values computed with the *eta_squared* function from the *effectsize* R package. Post hoc analyses on ANOVA effects were computed with the *emmeans* and *contrast* functions from the *emmeans* R package (*p*-values adjusted with Tukey’s method). Correlations were computed with the *spearmanr* (r_s_) or *pearsonr* (r_p_) function from the *stats* module of the Python *SciPy* package (v1.9.1), and two-sided t-tests were performed with the *ttest_ind* function from *SciPy.stats.* LASSO regression was implemented using the *Scikit-Learn* Python module (v1.0.2).

## Data & Code Availability

Data and analysis pipelines (code) used for this project will be made freely available on GitHub upon publication. The colorblind-friendly color schemes used in our figures were created using the toolbox available at https://davidmathlogic.com/colorblind/. Requests for further information should be directed to the Corresponding Author.

## Acknowledgements

This research was supported by a NIH T32 training grant (Maboudian).

Alzheimer’s Disease Neuroimaging Initiative: Data collection and sharing for this project was funded by ADNI (National Institutes of Health Grant U01 AG024904) and DOD ADNI (Department of Defense award number W81XWH-12-2-0012). ADNI is funded by the National Institute on Aging, the National Institute of Biomedical Imaging and Bioengineering, and through contributions from the following: AbbVie, Alzheimer’s Association; Alzheimer’s Drug Discovery Foundation; Araclon Biotech; BioClinica Inc.; Biogen; Bristol-Myers Squibb Company; CereSpir Inc.; Cogstate; Eisai Inc.; Elan Pharmaceuticals Inc.; Eli Lilly and Company; EuroImmun; F. Hoffmann-La Roche Ltd. and its affiliated company Genentech Inc.; Fujirebio; GE Healthcare; IXICO Ltd.; Janssen Alzheimer Immunotherapy Research & Development LLC.; Johnson & Johnson Pharmaceutical Research & Development LLC.; Lumosity; Lundbeck; Merck & Co Inc.; Meso Scale Diagnostics LLC.; NeuroRx Research; Neurotrack Technologies; Novartis Pharmaceuticals Corporation; Pfizer Inc.; Piramal Imaging; Servier; Takeda Pharmaceutical Company; and Transition Therapeutics. The Canadian Institutes of Health Research is providing funds to support ADNI clinical sites in Canada. Private sector contributions are facilitated by the Foundation for the National Institutes of Health (www.fnih.org). The grantee organization is the Northern California Institute for Research and Education, and the study is coordinated by the Alzheimer’s Therapeutic Research Institute at the University of Southern California. ADNI data are disseminated by the Laboratory for Neuro Imaging at the University of Southern California. Human Connectome Project (HCP): Young adult neuroimaging data were provided by the HCP, WU-Minn Consortium (Principal Investigators: David Van Essen and Kamil Ugurbil; NIH Grant 1U54-MH-091657) funded by the 16 NIH Institutes and Centers that support the NIH Blueprint for Neuroscience Research, and the McDonnell Center for Systems Neuroscience at Washington University.

## Author Contributions

S.A.M., W.J.J., and K.S.W. designed research. S.A.M., E.H.W., W.J.J., and K.S.W. performed research. S.A.M. analyzed data. S.A.M., W.J.J., and K.S.W. wrote the paper. All authors edited the paper and gave final approval before submission.

