## Supplementary Material for "Defining overlooked structures reveals new associations between cortex and cognition in aging and Alzheimer’s disease"

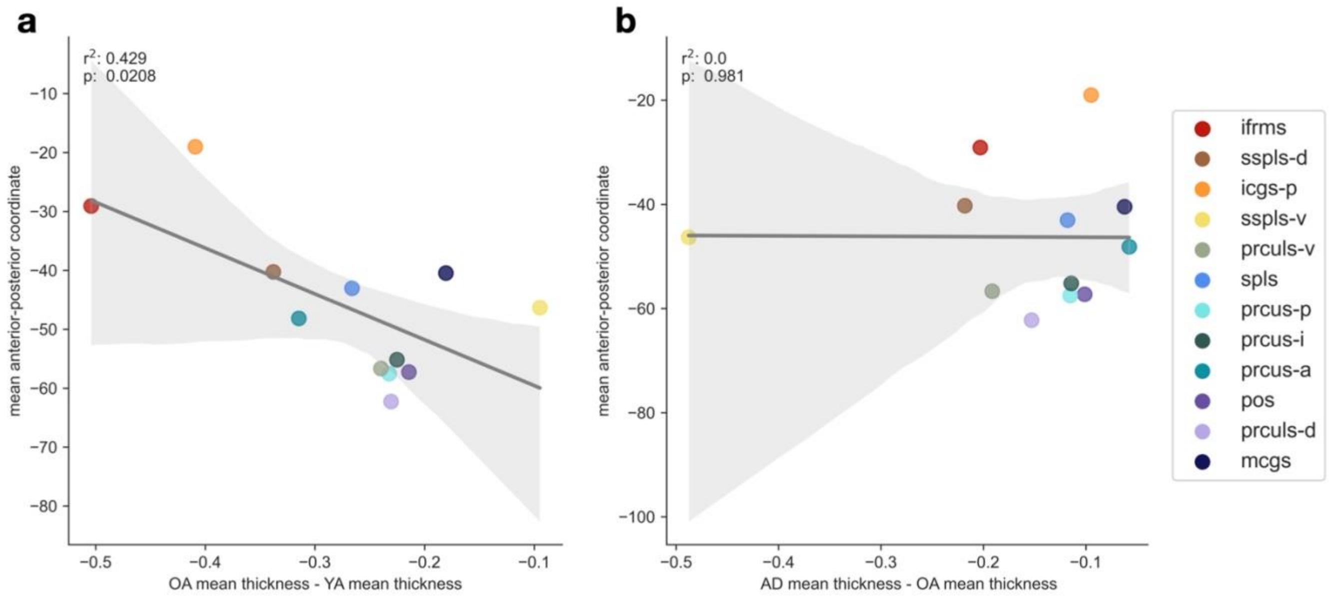


**Supplementary Figure 1. More anterior PMC sulci atrophy most in aging, but this anterior bias is absent in AD.**

**a** Mean anterior-posterior coordinate of each sulcus vs. the difference between the average thickness of the sulcus in OA and YA, demonstrating that more anterior sulci are more atrophied in OA. **b** Mean anterior-posterior coordinate of each sulcus vs. the difference between the average thickness of the sulcus in OA and AD, showing no relationship between anterior-posterior location and atrophy. Sulci colored based on legend (right).

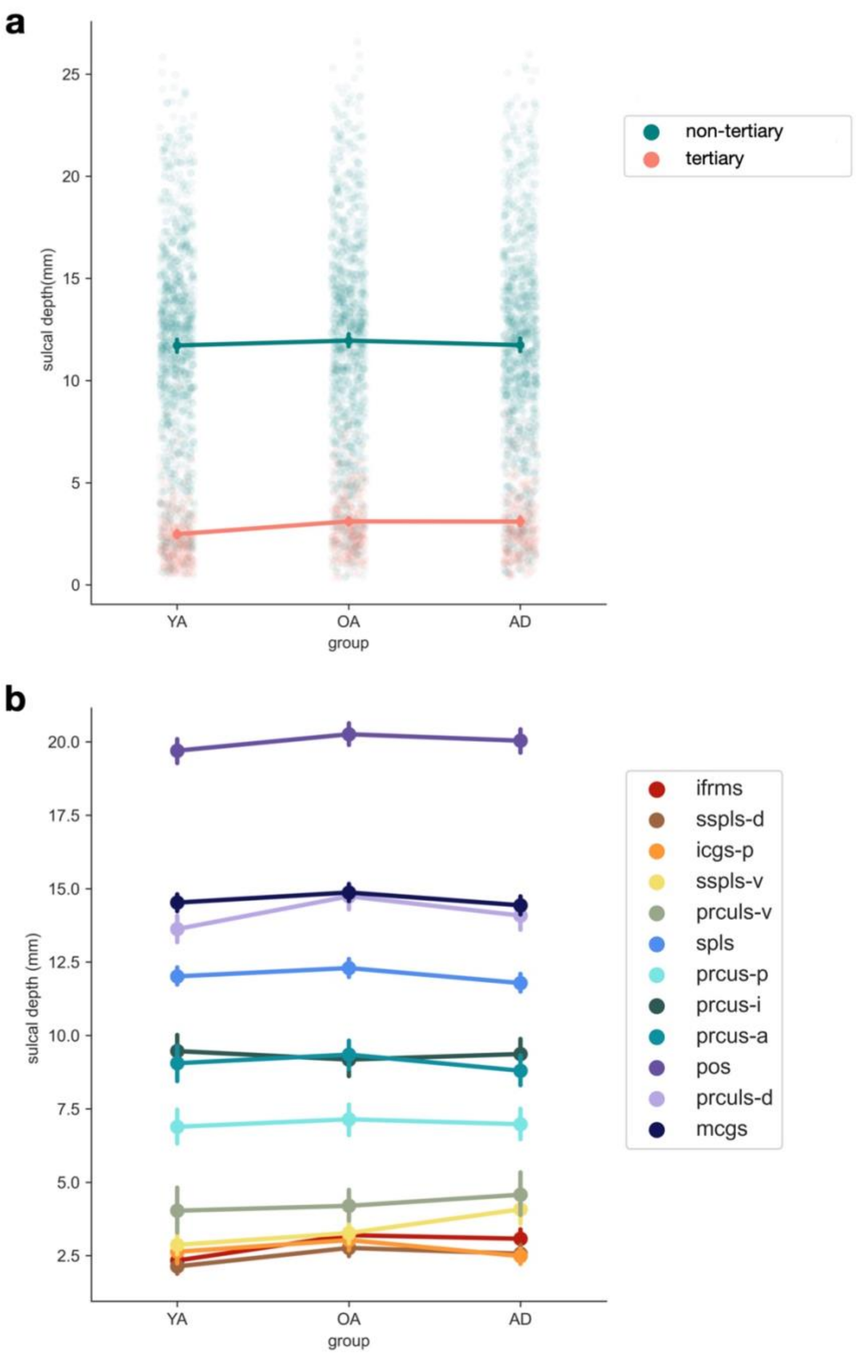


**Supplementary Figure 2. Sulcal depth comparisons across groups, by sulcal type or sulcus.**
**a** Sulcal depth (in mm) across groups (YA=younger adult, OA=cognitively normal older adult, AD=older adult with Alzheimer's disease) by sulcal type. A linear mixed effects (LME) model with group (YA, OA, AD), sulcal type (tertiary or non-tertiary), hemisphere (left or right) and their interactions as predictors shows a main effect of group ($p<0.05$), sulcal type ($p<0.0001$), and hemisphere ($p<0.0001$), and no significant interactions. The points represent the means, while vertical bars represent the bootstrap 95% CI. **b** Same as (**a**), but showing each sulcus separately, colored according to the legend. An LME model with group (YA, OA, AD), sulcus, hemisphere (left or right) and their interactions as predictors shows a main effect of group ($p<0.01$), sulcus ($p<0.0001$), and hemisphere ($p<0.0001$), as well as a group x sulcus interaction ($p<0.05$) and a sulcus x hemisphere interaction ($p<0.0001$).

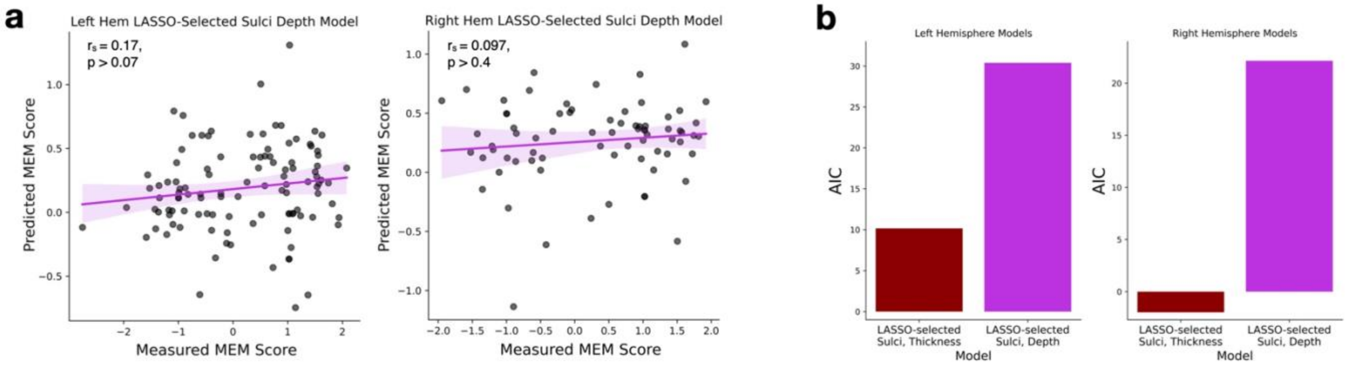


**Supplementary Figure 3. The depths of LASSO-selected sulci are not associated with ADNI-Mem scores.**

**a** Spearman's correlation ($r_S$) between actual ADNI Memory Composite (ADNI-MEM) scores and predicted scores from a linear regression model using LASSO-selected PMC sulcal depth to predict ADNI-MEM scores instead of LASSO-selected PMC sulcal thickness (as in Fig. 4), for both hemispheres. **b** Model comparison (AIC) of LASSO-selected sulci thickness and depth models (left: left hemisphere sulcal thickness models, right: right hemisphere models); ADNI-MEM thickness models have a substantially lower AIC in both cases.

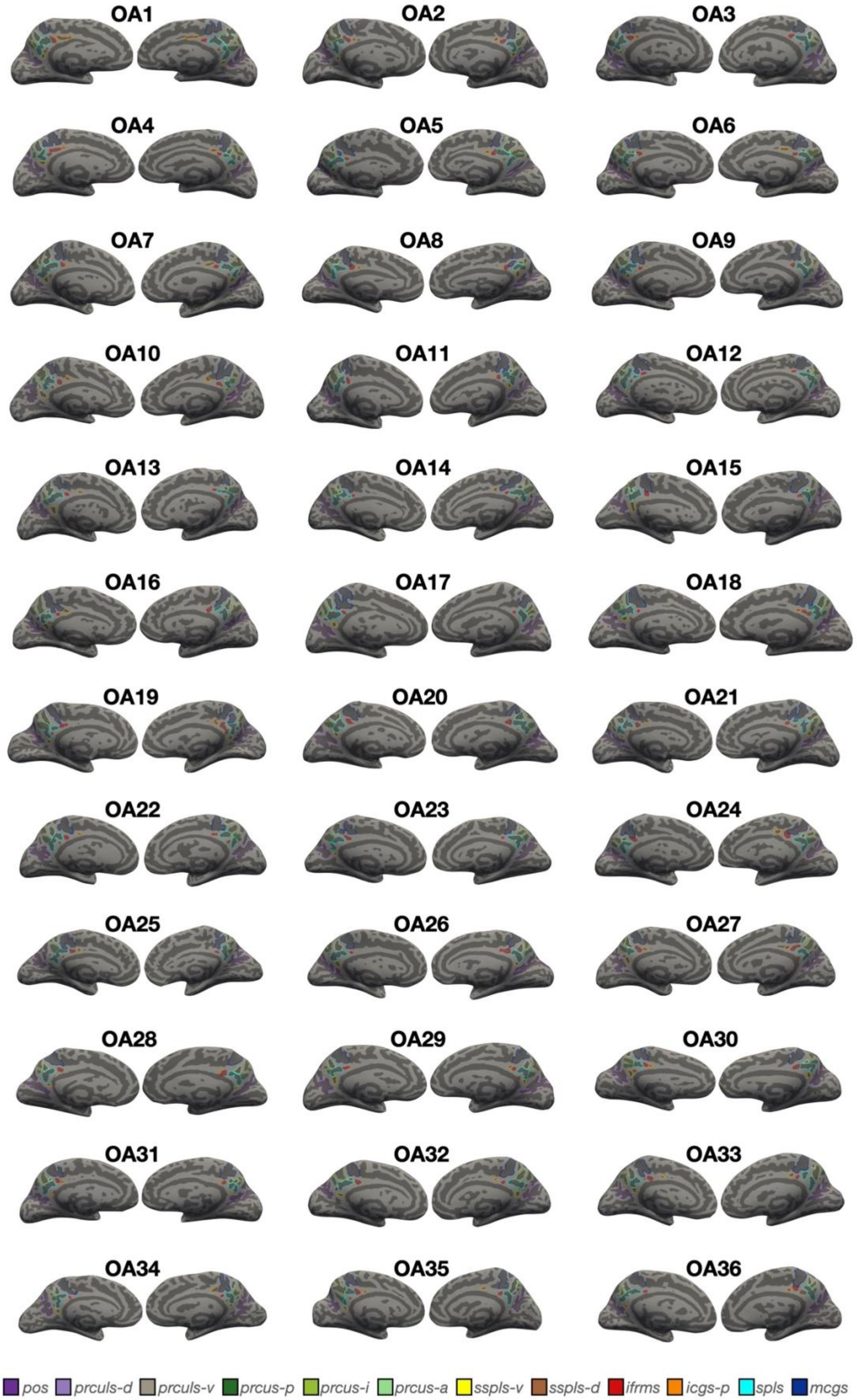
OA1
OA2
OA3
OA4
OA5
OA6
OA7
OA8
OA9
OA10
OA11
OA12
OA13
OA14
OA15
OA16
OA17
OA18
OA19
OA20
OA21
OA22
OA23
OA24
OA25
OA26
OA27
OA28
OA29
OA30
OA31
OA32
OA33
OA34
OA35
OA36
pos
prculs-d
prculs-v
prcus-p
prcus-i
prcus-a
sspls-v
sspls-d
ifrms
icgs-p
spls
mcgs

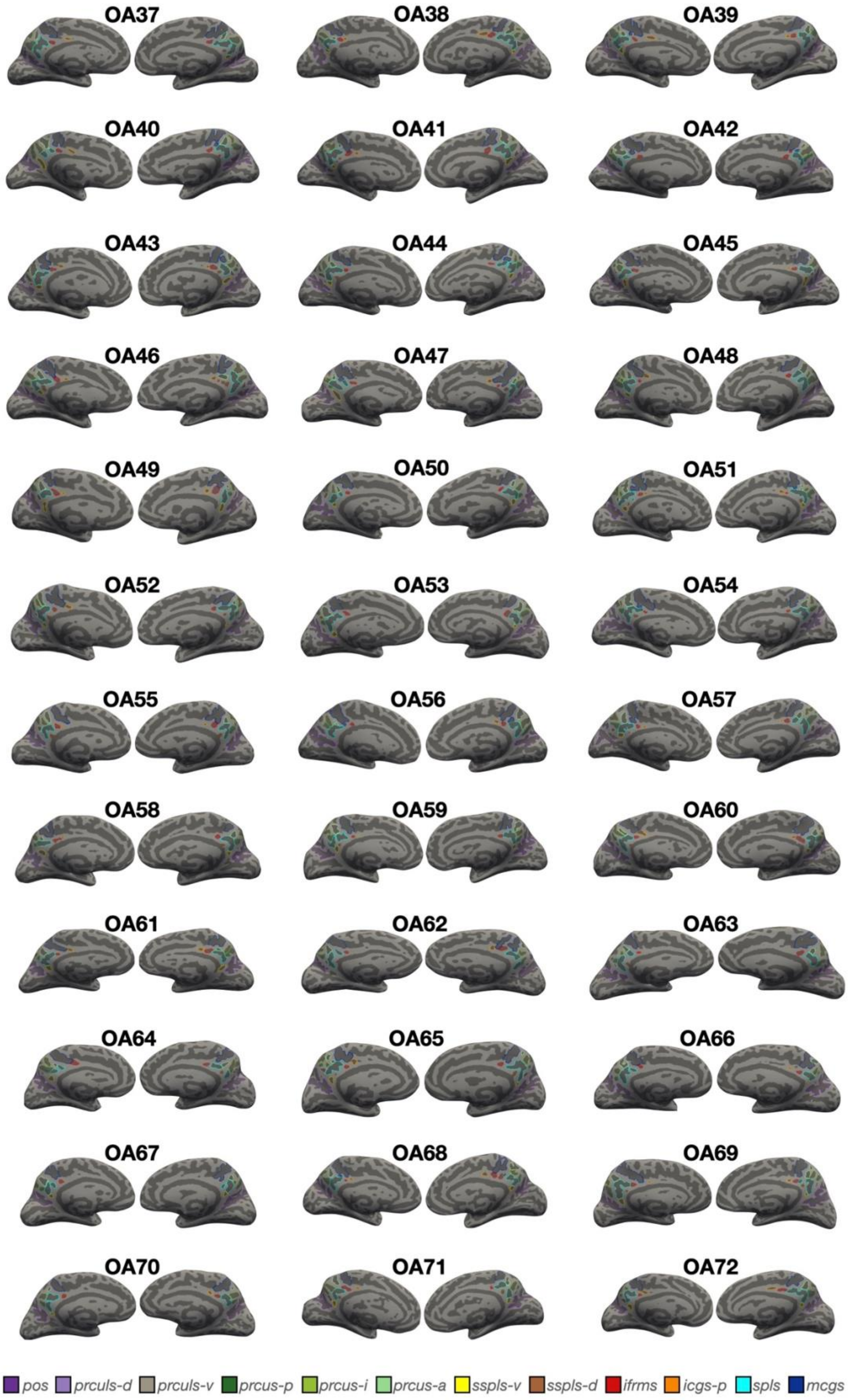
OA37
OA38
OA39
OA40
OA41
OA42
OA43
OA44
OA45
OA46
OA47
OA48
OA49
OA50
OA51
OA52
OA53
OA54
OA55
OA56
OA57
OA58
OA59
OA60
OA61
OA62
OA63
OA64
OA65
OA66
OA67
OA68
OA69
OA70
OA71
OA72
pos
prculs-d
prculs-v
prcus-p
prcus-i
prcus-a
sspls-v
sspls-d
ifrms
icgs-p
spls
mcgs

**Supplementary Figure 4. Manual PMC sulcal labels in every cognitively normal older adult (OA) participant.** Each sulcus is displayed on the left and right hemisphere inflated cortical surfaces in FreeSurfer 6.0.0, with label displayed as an outline according to the key at the top. Each hemisphere contains at least 8 sulci (from posterior to anterior): pos, prculs-d, prcus-p, prcus-i, prcus-a, spls, mcgs, and ifrms. An additional 4 sulci are variably present: prculs-v, sspls-v, sspls-d, icgs-p.

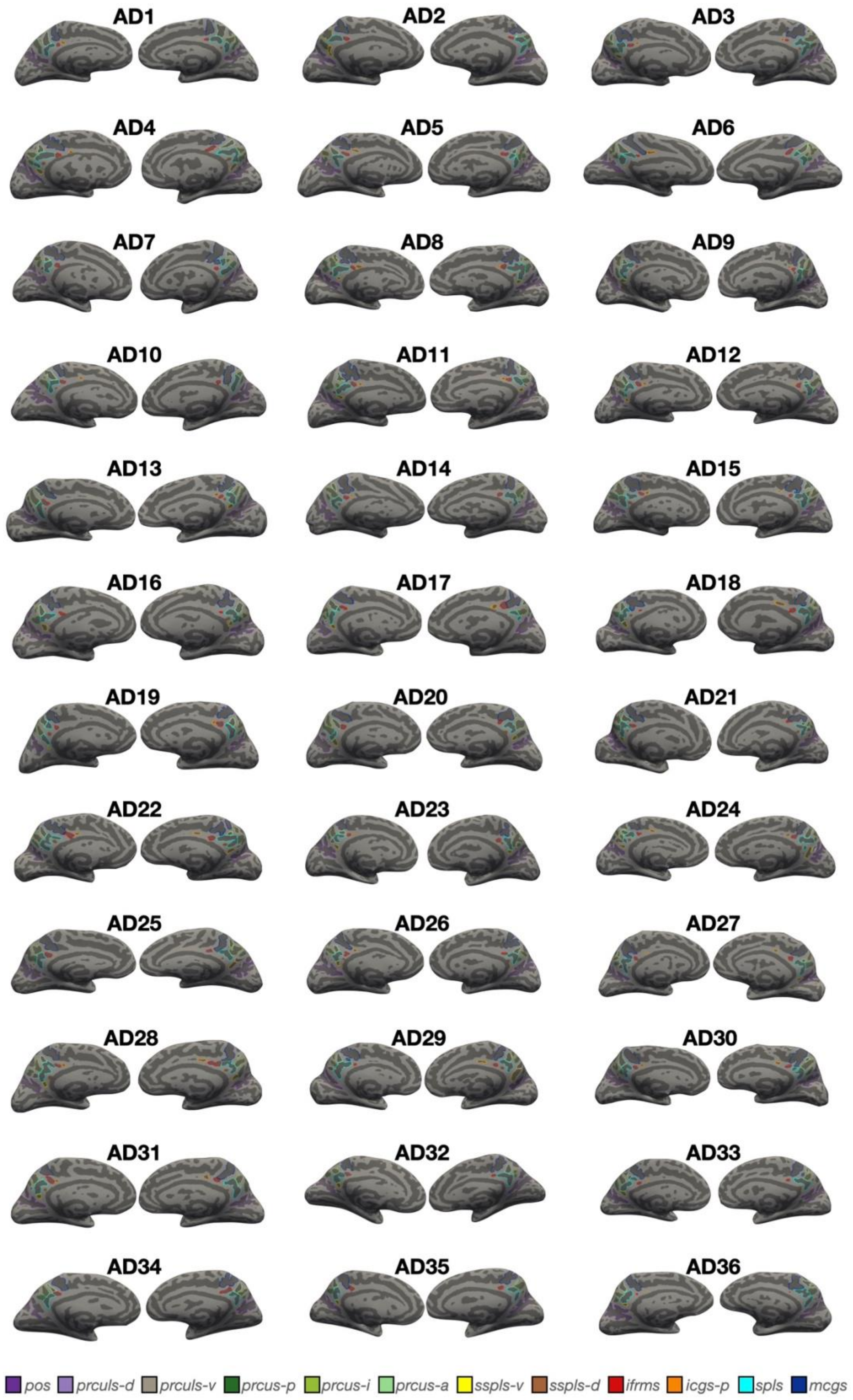
AD1
AD2
AD3
AD4
AD5
AD6
AD7
AD8
AD9
AD10
AD11
AD12
AD13
AD14
AD15
AD16
AD17
AD18
AD19
AD20
AD21
AD22
AD23
AD24
AD25
AD26
AD27
AD28
AD29
AD30
AD31
AD32
AD33
AD34
AD35
AD36
pos
prculs-d
prculs-v
prcus-p
prcus-i
prcus-a
sspls-v
sspls-d
ifrms
icgs-p
spls
mcgs

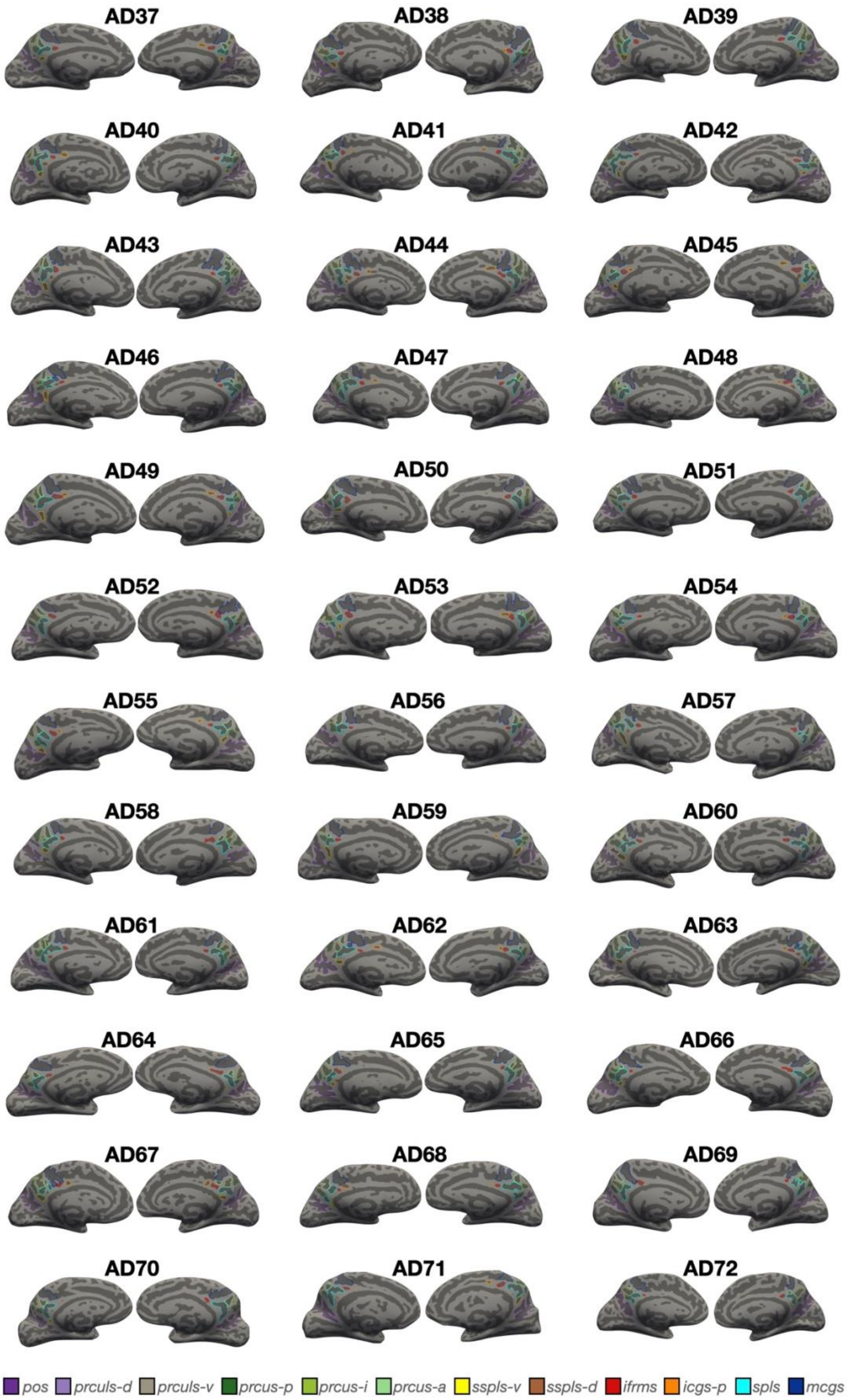
AD37
AD38
AD39
AD40
AD41
AD42
AD43
AD44
AD45
AD46
AD47
AD48
AD49
AD50
AD51
AD52
AD53
AD54
AD55
AD56
AD57
AD58
AD59
AD60
AD61
AD62
AD63
AD64
AD65
AD66
AD67
AD68
AD69
AD70
AD71
AD72
pos
prculs-d
prculs-v
prcus-p
prcus-i
prcus-a
sspls-v
sspls-d
ifrms
icgs-p
spls
mcgs

**Supplementary Figure 5. Manual PMC sulcal labels in every older adult participant with Alzheimer's disease (AD).** Each sulcus is displayed on the left and right hemisphere inflated cortical surfaces in FreeSurfer 6.0.0, with label displayed as an outline according to the key at the top. Each hemisphere contains at least 8 sulci (from posterior to anterior): pos, prculs-d, prcus-p, prcus-i, prcus-a, spls, mcgs, and ifrms. An additional 4 sulci are variably present: prculs-v, sspls-v, sspls-d, icgs-p.

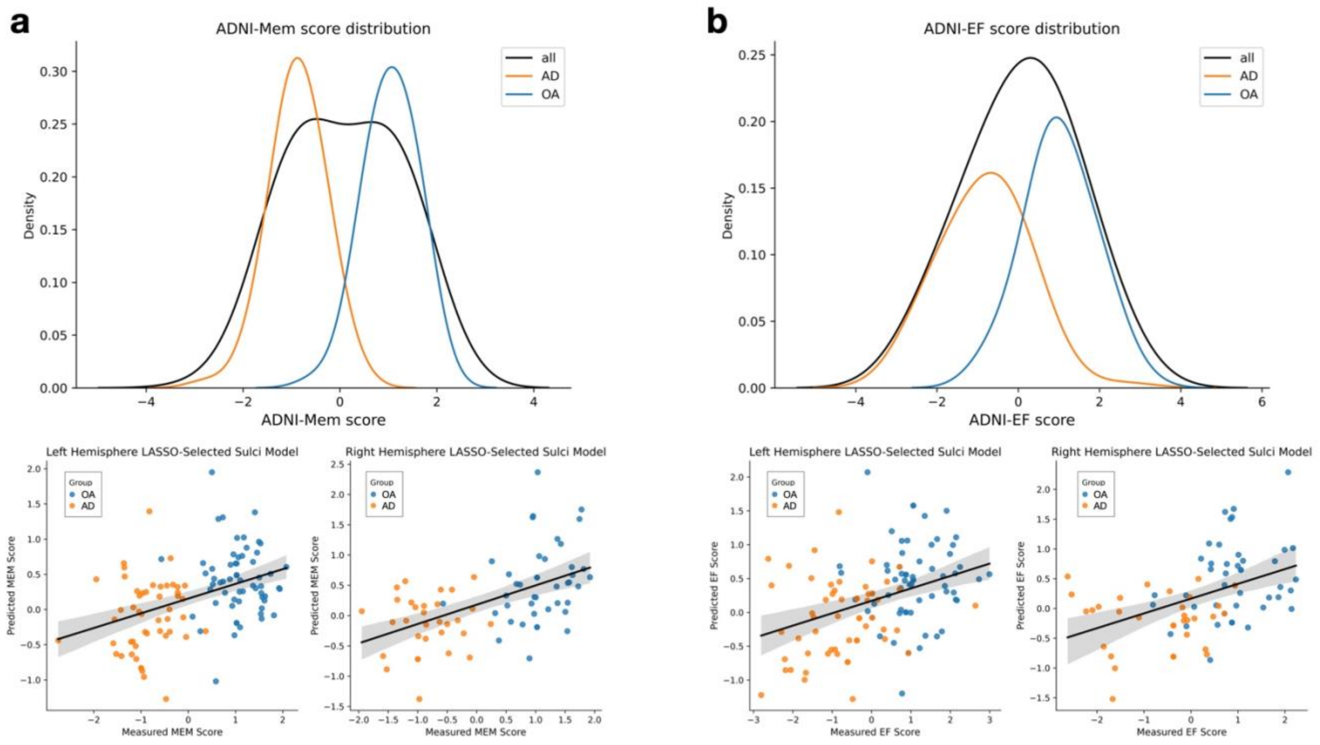


**Supplementary Figure 6. Distribution of cognitive scores across groups.**
**a** Distribution of ADNI Memory Composite (ADNI-Mem) scores by group (top), and model prediction accuracy of models in Fig 5 split by group (bottom). **b** Same as (**a**), but for Executive Function Composite (ADNI-EF) scores.

**a**

| Sulcal type | Group comparison | Estimate | t-ratio | p-value |
|---|---|---|---|---|
| Non-tertiary | OA-YA | -0.234 | -9.412 | **<.0001** |
| | AD-OA | -0.112 | -4.500 | **<.0001** |
| Tertiary | OA-YA | -0.355 | -11.650 | **<.0001** |
| | AD-OA | -0.243 | -7.831 | **<.0001** |

**b**

| Sulcus | Group comparison | Estimate | t-ratio | p-value |
|---|---|---|---|---|
| icgs-p | OA-YA | -0.3946 | -8.437 | **<.0001** |
| | AD-OA | -0.0992 | -1.987 | 0.118 |
| ifrms | OA-YA | -0.5031 | -13.017 | **<.0001** |
| | AD-OA | -0.2041 | -5.275 | **<.0001** |
| sspls-d | OA-YA | -0.3596 | -6.860 | **<.0001** |
| | AD-OA | -0.1975 | -3.624 | **0.001** |
| sspls-v | OA-YA | -0.0964 | -2.052 | 0.10 |
| | AD-OA | -0.4956 | -10.354 | **<.0001** |
| mcgs | OA-YA | -0.1803 | -4.670 | **<.0001** |
| | AD-OA | -0.0622 | -1.612 | 0.24 |
| pos | OA-YA | -0.2143 | -5.552 | **<.0001** |
| | AD-OA | -0.1013 | -2.624 | **0.025** |
| prculs-d | OA-YA | -0.2307 | -5.976 | **<.0001** |
| | AD-OA | -0.1531 | -3.967 | **<.001** |
| prculs-v | OA-YA | -0.2578 | -4.897 | **<.0001** |
| | AD-OA | -0.1839 | -3.564 | **0.0013** |
| prcus-p | OA-YA | -0.2324 | -6.021 | **<.0001** |
| | AD-OA | -0.1154 | -2.989 | **0.009** |
| prcus-i | OA-YA | -0.2252 | -5.835 | **<.0001** |
| | AD-OA | -0.1143 | -2.962 | **0.0095** |
| prcus-a | OA-YA | -0.3147 | -8.155 | **<.0001** |
| | AD-OA | -0.0578 | -1.494 | 0.30 |
| spls | OA-YA | -0.2663 | -6.899 | **<.0001** |
| | AD-OA | -0.1179 | -3.056 | **0.007** |

**Supplementary Table 1. Results of post-hoc tests for linear mixed effects (LME) models of cortical thickness. a** Results from post-hoc comparisons for the model comparing cortical thickness across groups by sulcal type (Fig. 3A). Each pairwise group comparison is significant for each sulcal type category separately, but group differences are greater among tertiary sulci. **b** Results from post-hoc comparisons for the model comparing the cortical thickness of every PMC sulcus (Fig. 3B), calculated with *emmeans* in R. Pairwise group comparisons are significant (Tukey's-adjusted $p < 0.05$) for every sulcus except icgs-p ($p$=0.12), mcgs ($p$=0.24) and prcus-a ($p$=0.30) for the AD-OA comparison, and sspls-v for the OA-YA comparison ($p$=0.10). The strongest effects are seen in the ifrms for the OA-YA comparison and the sspls-v for the AD-OA comparison.

| **Group** | **Age:** range (μ, sd) | **Sex:** %F | **Years of Edu:** μ (sd) | **Aβ Status:** % Aβ+ |
|---|---|---|---|---|
| YA | 22-36 (29.06, 3.57) | 50.0 | N/A | N/A |
| OA | 65-90 (76.01, 6.32) | 52.8 | 16.47 (2.25) | 51% |
| AD | 65-89 (76.15, 5.89) | 48.6 | 15.74 (2.32) | 100% |

**Supplementary Table 2. Participant characteristics.** This table illustrates participant characteristics of our samples from the Human Connectome Project (HCP-YA) and the Alzheimer's Disease Neuroimaging Initiative (ADNI) databases. Amyloid (Aβ) status was only available in the older adult groups (OA and AD), and years of education (Edu) is only reported for older adult groups because only older adults are considered in cognition analyses.

| Manufacturer | Field Strength | TE | TR | Pixel Spacing X | Pixel Spacing Y | Slice Thickness | count |
|---|---|---|---|---|---|---|---|
| SIEMENS | 3.0 | 2.980 | 2300.0000 | 1.00000 | 1.00000 | 1.0 | 32 |
| SIEMENS | 3.0 | 2.980 | 2300.0000 | 1.00000 | 1.00000 | 1.2 | 15 |
| SIEMENS | 3.0 | 2.950 | 2300.0000 | 1.05469 | 1.05469 | 1.2 | 11 |
| Philips Medical Systems | 3.0 | 3.160 | 6.8054 | 1.00000 | 1.00000 | 1.2 | 3 |
| Philips Medical Systems | 3.0 | 2.942 | 6.5175 | 1.00000 | 1.00000 | 1.0 | 2 |
| SIEMENS | 3.0 | 2.960 | 2300.0000 | 1.00000 | 1.00000 | 1.0 | 2 |
| GE MEDICAL SYSTEMS | 1.5 | 3.800 | 8.5920 | 0.93750 | 0.93750 | 1.2 | 1 |
| Philips Medical Systems | 3.0 | 2.935 | 6.4461 | 1.00000 | 1.00000 | 1.0 | 1 |
| Philips Medical Systems | 3.0 | 2.944 | 6.5362 | 1.00000 | 1.00000 | 1.0 | 1 |
| Philips Medical Systems | 3.0 | 2.944 | 6.5364 | 1.00000 | 1.00000 | 1.0 | 1 |
| Philips Medical Systems | 3.0 | 3.133 | 6.7702 | 1.00000 | 1.00000 | 1.2 | 1 |
| Philips Medical Systems | 3.0 | 3.137 | 6.7669 | 1.00000 | 1.00000 | 1.2 | 1 |
| Philips Medical Systems | 3.0 | 3.157 | 6.7805 | 1.00000 | 1.00000 | 1.2 | 1 |

**Supplementary Table 3. Scanning parameters of the scans from cognitively normal older adult participants**. This table illustrates the different scanning parameters used for each of the cognitively normal older adult participants from the Alzheimer's Disease Neuroimaging Initiative (ADNI) online database (http://adni.loni.usc.edu).

| Manufacturer | Field Strength | TE | TR | Pixel Spacing X | Pixel Spacing Y | Slice Thickness | count |
|---|---|---|---|---|---|---|---|
| SIEMENS | 3.0 | 2.980 | 2300 | 1.00000 | 1.00000 | 1.2 | 33 |
| SIEMENS | 3.0 | 2.980 | 2300 | 1.00000 | 1.00000 | 1.0 | 13 |
| Philips Medical Systems | 3.0 | 3.133 | 6.7702 | 1.00000 | 1.00000 | 1.2 | 5 |
| Philips Medical Systems | 3.0 | 3.157 | 6.7805 | 1.00000 | 1.00000 | 1.2 | 3 |
| SIEMENS | 3.0 | 2.950 | 2300 | 1.05469 | 1.05469 | 1.2 | 2 |
| Philips Medical Systems | 3.0 | 3.137 | 6.7671 | 1.00000 | 1.00000 | 1.2 | 2 |
| Philips Medical Systems | 3.0 | 3.136 | 6.7742 | 1.00000 | 1.00000 | 1.2 | 1 |
| Philips Medical Systems | 3.0 | 3.160 | 6.8054 | 1.00000 | 1.00000 | 1.2 | 1 |
| Philips Medical Systems | 3.0 | 3.153 | 6.8011 | 1.00000 | 1.00000 | 1.2 | 1 |
| Philips Medical Systems | 3.0 | 3.137 | 6.7674 | 1.00000 | 1.00000 | 1.2 | 1 |
| Philips Healthcare | 3.0 | 2.944 | 6.5362 | 1.00000 | 1.00000 | 1.0 | 1 |
| Philips Healthcare | 3.0 | 2.944 | 6.5364 | 1.00000 | 1.00000 | 1.0 | 1 |
| Philips Medical Systems | 3.0 | 3.133 | 6.7710 | 1.00000 | 1.00000 | 1.2 | 1 |
| Philips Medical Systems | 3.0 | 3.112 | 6.7283 | 1.00000 | 1.00000 | 1.2 | 1 |
| Philips Medical Systems | 3.0 | 2.944 | 6.5364 | 1.00000 | 1.00000 | 1.0 | 1 |
| Philips Medical Systems | 3.0 | 2.944 | 6.5362 | 1.00000 | 1.00000 | 1.0 | 1 |
| Philips Medical Systems | 3.0 | 2.942 | 6.5160 | 1.00000 | 1.00000 | 1.0 | 1 |
| Philips Medical Systems | 3.0 | 2.940 | 6.5126 | 1.00000 | 1.00000 | 1.0 | 1 |
| Philips Medical Systems | 3.0 | 2.928 | 6.5414 | 1.00000 | 1.00000 | 1.0 | 1 |

| | | | | | | | |
|---|---|---|---|---|---|---|---|
| Philips Medical Systems | 3.0 | 3.136 | 6.7740 | 1.00000 | 1.00000 | 1.2 | 1 |

**Supplementary Table 4. Scanning parameters of the scans from older adult participants with Alzheimer's disease.** This table illustrates the different scanning parameters used for each of the older adult participants with Alzheimer's disease from the Alzheimer's Disease Neuroimaging Initiative (ADNI) online database (http://adni.loni.usc.edu).
